# pLM-BLAST – distant homology detection based on direct comparison of sequence representations from protein language models

**DOI:** 10.1101/2022.11.24.517862

**Authors:** Kamil Kaminski, Jan Ludwiczak, Kamil Pawlicki, Vikram Alva, Stanislaw Dunin-Horkawicz

**Author notes:** Prescient Design, Genentech Research & Early Development, Roche Group, Grenzacherstrasse 124, 4070 Basel, Switzerland (Jan Ludwiczak).

## Abstract

**Motivation:** The detection of homology through sequence comparison is a typical first step in the study of protein function and evolution. In this work, we explore the applicability of protein language models to this task.

**Results:** We introduce pLM-BLAST, a tool inspired by BLAST, that detects distant homology by comparing single-sequence representations (embeddings) derived from a protein language model, ProtT5. Our benchmarks reveal that pLM-BLAST maintains a level of accuracy on par with HHsearch for both highly similar sequences (with over 50% identity) and markedly divergent sequences (with less than 30% identity), while being significantly faster. Additionally, pLM-BLAST stands out among other embedding-based tools due to its ability to compute local alignments. We show that these local alignments, produced by pLM-BLAST, often connect highly divergent proteins, thereby highlighting its potential to uncover previously undiscovered homologous relationships and improve protein annotation.

**Availability and Implementation:** pLM-BLAST is accessible via the MPI Bioinformatics Toolkit as a web server for searching precomputed databases (https://toolkit.tuebingen.mpg.de/tools/plmblast). It is also available as a standalone tool for building custom databases and performing batch searches (https://github.com/labstructbioinf/pLM-BLAST).

## Introduction

Homology, i.e., descent from a common ancestor, and its inference are fundamental to comparative, evolutionary, and molecular biology. In the case of proteins, statistically significant local or global sequence similarity is accepted as the primary marker for inferring homology. When the similarity between the protein sequences being compared is high (>=30%), homology can be readily detected using methods based on sequence-to-sequence and sequence-to-profile comparisons such as BLAST or PSI-BLAST (Altschul *et al*., 1997). However, when similarity is low (<30%), methods based on profile HMMs, such as HMMER and HHsearch, are currently our best tools to infer homology (Steinegger *et al*., 2019; Eddy, 2011).

In the case of highly distant evolutionary relationships, sequences may have diverged to the point where we can no longer detect their relatedness. Because structures diverge much more slowly than sequences, their similarity is often used to infer homology in such cases. In fact, owing to the recent revolution in structure prediction (Lin *et al*., 2022; Jumper *et al*., 2021; Li *et al*., 2022), predicted structures are available for a large proportion of known proteins, and structural similarity is increasingly being used to infer homology. However, similar structures, particularly at the domain and sub-domain levels, may have evolved convergently due to the limited number of structural solutions available to a folded polypeptide chain, and thus structural similarity is often not conclusive evidence of common ancestry. To address this issue, several deep learning-based methods have sought to detect weak sequence signals between highly divergent proteins in recent years (Zheng *et al*., 2019; Gao and Skolnick, 2021; Li *et al*., 2017). The most promising of these methods, such as knnProtT5 (Schütze *et al*., 2022), EBA (Pantolini *et al*., 2022), TM-Vec (Hamamsy *et al*., 2022), DeepBLAST (Morton *et al*., 2020), and DEDAL (Llinares-López *et al*., 2023), rely on single-sequence representations obtained from protein language models (pLMs) (Bepler and Berger, 2021). pLMs are neural networks trained in a self-supervised manner on a large number of natural protein sequences, i.e., with tasks such as guessing masked residues based on contextual information (Elnaggar *et al*., 2021; Lin *et al*., 2022). Once trained, pLMs can be used to rapidly compute the aforementioned representations (also referred to as embeddings), i.e., numerical descriptors of protein sequences that place them in the context of the total knowledge collected by the network. Because of their high information content, sequence embeddings have been successfully applied to many other tasks, including prediction of tertiary structures (Lin *et al*., 2022), transmembrane segments (Bernhofer and Rost, 2022), and signal peptides (Teufel *et al*., 2022).

In this article, we describe pLM-BLAST, a new tool for detecting local homology between protein sequences that combines pLM representations with a local similarity detection algorithm inspired by BLAST (Altschul *et al*., 1997). In contrast to TM-vec and DEDAL, pLM-BLAST is based on an unsupervised approach that does not require training of a specialized deep-learning model or defining positive labels based on structurally similar or homologous protein pairs. Furthermore, unlike methods such as TM-vec and EBA, which only provide global alignments, pLM-BLAST has the ability to compute both local and global alignments, which is essential for determining distant homology relationships that may depend on the conservation of short subdomain fragments (Alva *et al*., 2015; Kolodny, 2021). pLM-BLAST is based on an implementation of a modified Smith-Waterman algorithm, in which the substitution matrix is generated directly from the raw pLM representations of the two sequences to be compared. In this work, we used the pLM ProtT5 (Elnaggar *et al*., 2021), but this approach can be used in conjunction with other pLMs to generate local and global alignments. In benchmarks using domain pairs from the ECOD protein classification database (Cheng *et al*., 2014), pLM-BLAST demonstrated its suitability for fast pairwise comparisons and database searches to detect distant homology and produce highly precise alignments.

## Methods

pLM-BLAST extends the concept of BLAST by replacing invariant substitution matrices, such as BLOSUM62 (Henikoff and Henikoff, 1992), with per-residue similarities between protein embeddings. Consequently, the similarity between a given pair of residues is entirely context-dependent. Such sequence context information has been shown to significantly improve the sensitivity of sequence search methods and enable the detection of non-trivial conserved patterns (Biegert and Söding, 2009; Remmert *et al*., 2011). Below is the workflow of the method, which is also shown in Figure 1.

**Figure 1.**
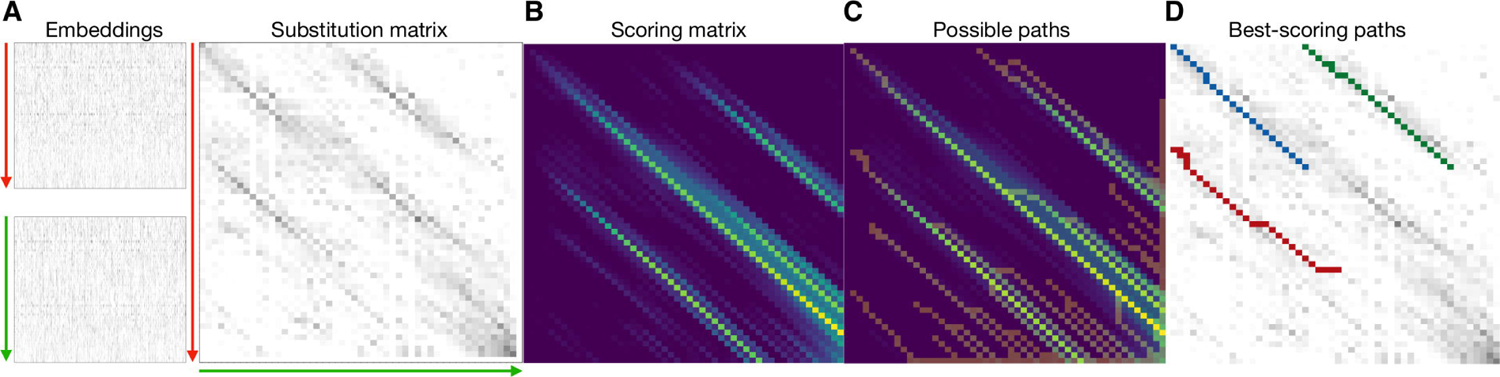
**(A)** Embeddings for the two sequences to be aligned. The y-axis represents the sequence dimension from N to C-terminus, and the x-axis represents the embedding dimension (which is a 1024-element vector in the case of the ProtT5 protein language model). The substitution matrix is computed by matrix multiplication of both embeddings, after each residue has been normalized. **(B)** The scoring matrix is computed to detect continuous high-scoring regions in the substitution matrix. **(C)** A modified traceback procedure is used to traverse the scoring matrix to extract potentially matching regions, called paths. **(D)** The possible paths are mapped back onto the substitution matrix, and a moving window of defined length is used to extract high-scoring sub-paths, i.e., local alignments.

### Substitution matrix

The embedding of a sequence *seq_1_*., is given by a matrix of size *n x m*, where *n* and *m* denote the number of residues and the size of the embedding, respectively. To obtain standardized and comparable results, the values for each residue embedding are normalized by dividing them by the Euclidean norm of the row

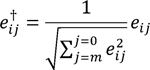

where *e_ij_* denotes the *E_l_* matrix element in the i-th row and j-th column, and *†* is the sequences and is calculated as normalization operator. In pLM-BLAST, a substitution matrix *S_lk_* (Figure 1A) for two sequences *seq_l_* and *seq_k_* is calculated as

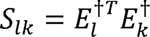

The element *S_ij_* of the matrix *S_lk_* is in the interval (−1,1) and denotes the embedding similarity of the i-th residue of *seq_L_* and the j-th residue of *seq_K_*. This is equivalent to calculating the cosine similarity of each pair of residues from *seq_L_* and *seq_K_* and placing them in a two-dimensional array indexed by the residues of each sequence.

#### Scoring matrix

To obtain local alignments, a scheme adapted from the Smith-Waterman algorithm is used to create a scoring matrix (Figure 1B) H_LK_ ∋ {h_ij_} of size (|L~ + 1) × (|K~ + 1), where the first row and column are filled with zeros and the rest of the matrix is filled using the following procedure

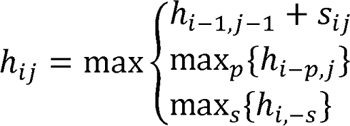

Unlike the original Smith-Waterman algorithm, pLM-BLAST does not use gap penalties because the _.i}_ values for dissimilar regions are negative or close to zero due to the cosine similarity property. In addition, the above equation does not truncate the values to zero, which results in a more severe penalty for dissimilar regions, thus reducing the total number of potential alignments.

#### Traceback procedure

To identify possible matching regions for a given pair of sequences, a traceback procedure is used to traverse the scoring matrix (Figure 1C). Unlike the SW algorithm, we don’t start with the highest value in the scoring matrix, but traverse from all sequence boundaries. Thus, in our approach, the traceback procedure does not produce a single alignment, but multiple candidates for possible alignments. For this purpose, possible paths are constructed, starting from the right and bottom edges of the matrix *H*. The path *p_n_* is a set of coordinates from the matrix *H_LK_*, where n ∋ N and N equals |L~ + |K~.

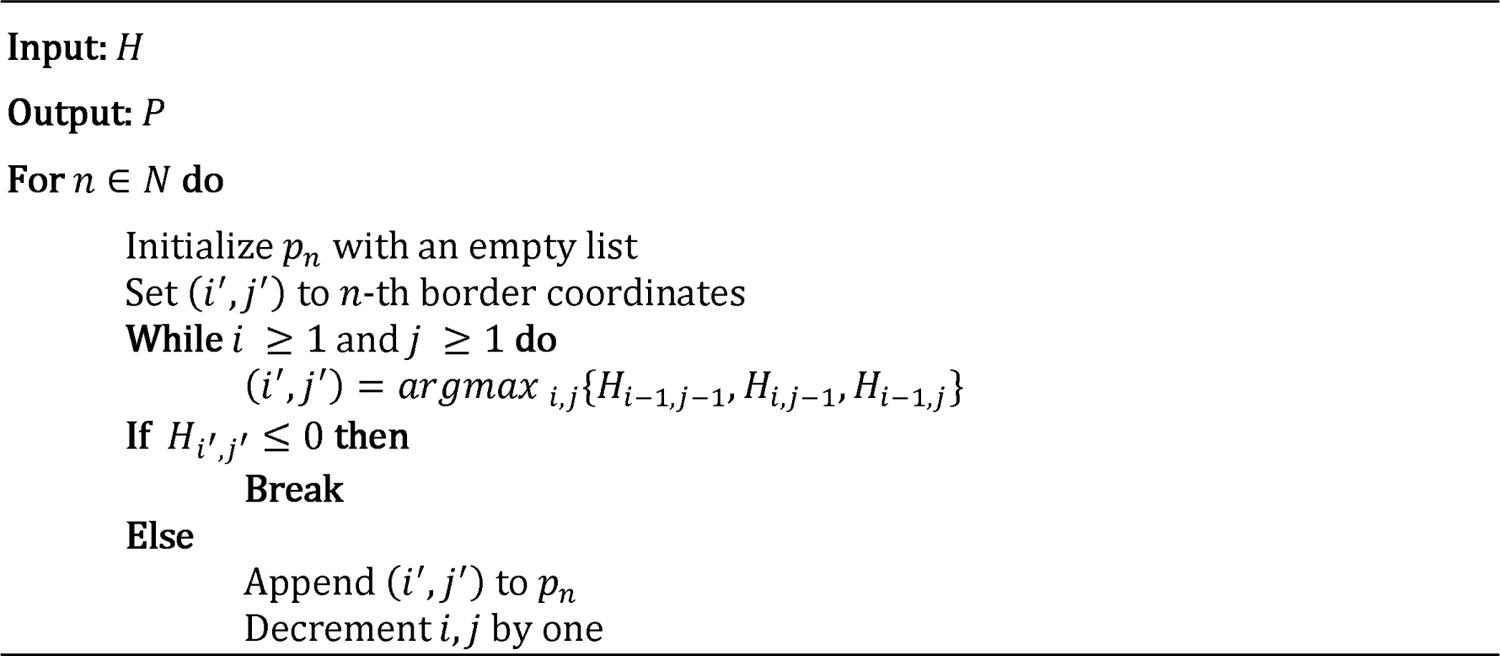

#### Local alignment

Each path generated by the traceback algorithm is scanned for the presence of high-scoring subpaths (i.e., local alignments) using the values of the corresponding region in the substitution matrix. To this end, a moving average with a defined fixed window length is applied to each path *p_n_* and subpaths whose average score is above the sigma threshold are captured. By default, the threshold is set to 2 sigmas, which is defined as 2 times the average standard deviation of the substitution matrix. Our tests showed that the standard deviation of the substitution matrix tends to be around 0.075, while the path score for meaningful alignments is an order of magnitude higher. However, increasing (>2) or decreasing (<2) the standard deviation cutoff makes the algorithm more strict or more permissive, respectively.

#### Global alignment

In addition to local alignments, pLM-BLAST can also perform global alignments using the Needleman-Wunsch algorithm (Needleman and Wunsch, 1970). The original algorithm remains the same except that the similarity matrix is replaced by the embedding-based substitution matrix S and the gap penalty is set to zero.

#### Homology benchmark

To evaluate the ability of pLM-BLAST and other methods to detect homology between protein sequences, we constructed two benchmark sets based on the ECOD database (Cheng *et al*., 2014): first, “hard”, which focuses on difficult cases where the similarity between sequences does not exceed 30%; and second, “easy”, where we test the methods in detecting homology between sequences with 50% to 70% similarity.

To construct the “hard” set, the sequences of all ECOD domains (version 20220912), excluding nested domains, with a length of 50-600 amino acids were obtained using the *localpdb* ECOD plugin (Ludwiczak *et al*., 2022) and clustered using MMSeqs2 with a sequence identity cutoff of 30% (Steinegger and Söding, 2017). The longest sequences were selected as cluster representatives and those belonging to H-groups with less than five members were removed. The final benchmark set was created by randomly selecting five domains from 300 H-groups, i.e., ECOD levels that group together homologous domains with common ancestry. This procedure was designed to select domains from different T-groups; a T-group is a sublevel that groups together homologous domains within an H-group that are topologically similar. As a result, we obtained a set of 1500 ECOD domains from 300 H-groups, none of which shared more than 30% sequence identity.

For each ECOD domain in the benchmark set, an HMM profile and sequence embedding were calculated using three iterations of HHblits over the UniRef30 database with default settings (Steinegger *et al*., 2019) and ProtT5 (prot_t5_xl_half_uniref50-enc) (Elnaggar *et al*., 2021), respectively. The embeddings were compared all-against-all with pLM-BLAST (in local and global mode), TM-Vec, and EBA, while the HMM profiles were compared with HHsearch. Matches between domains of the same H-group were considered true positives, matches between different H-groups belonging to the same X-group (an ECOD classification level that groups H-groups that may be homologous) were considered neutral, and finally matches between different X-groups were considered false positives. For the calculation of precision-recall curves (Figure 2), we filled missing data (i.e., pairs for which a given method did not provide a score) with zeros, assuming that the lack of prediction can be considered a negative prediction. In Figure 2, the regions of the curves resulting from such a completion procedure have been omitted for clarity (the complete curves are available as Supplementary Figure 1). We used the HHsearch match probability, the EBA max score, and the raw pLM-BLAST and TM-Vec scores to calculate the plots.

**Figure 2.**
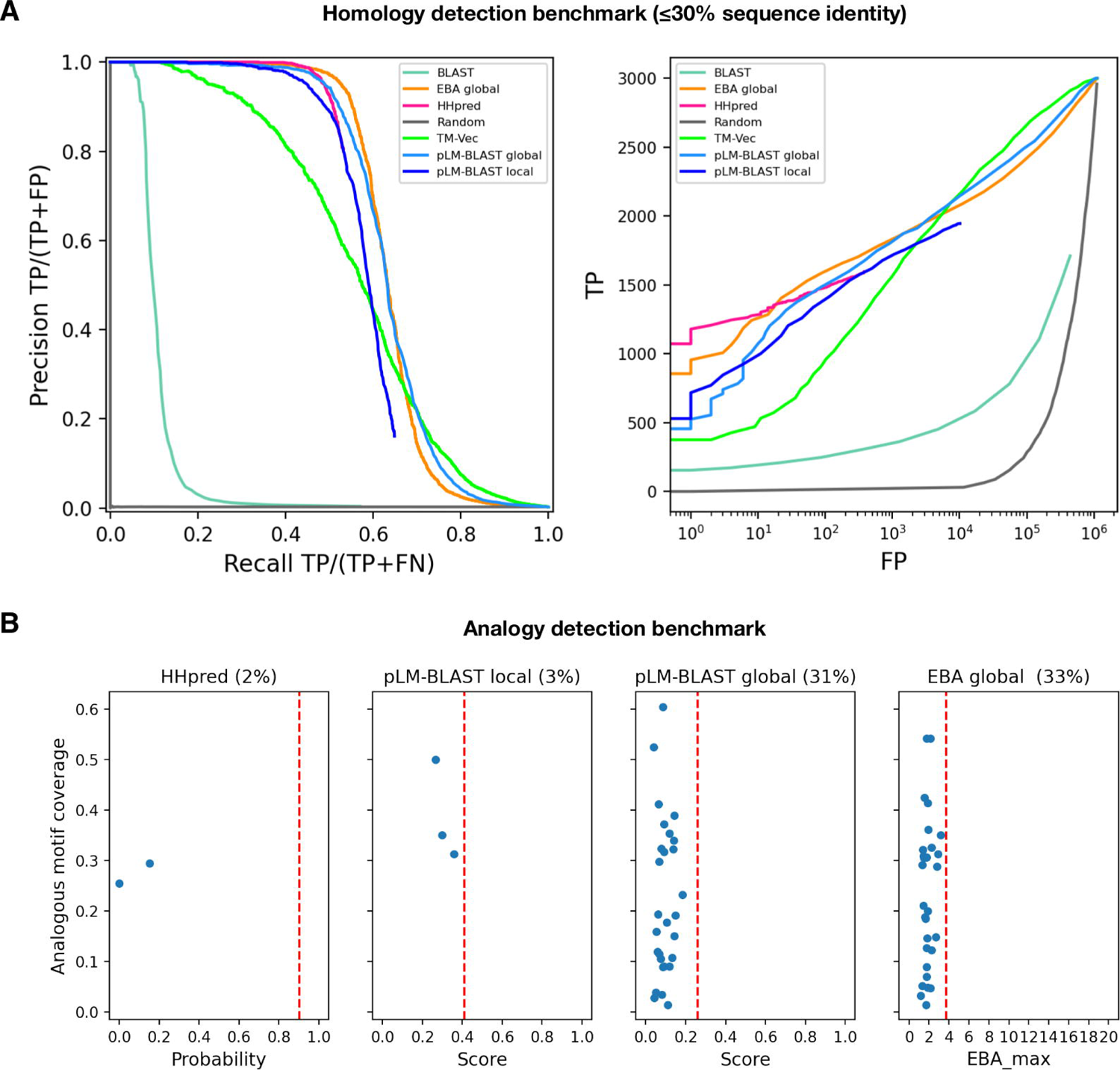
Performance of pLM-BLAST and other methods in detecting homology and analogy between protein sequences. **(A)** Benchmark for the task of reconstructing H-groups as defined in the ECOD database. The left panel shows the precision-recall curve and the right panel shows the number of true and false positives. Please note that the false positives are plotted on a logarithmic scale. **(B)**A comparison of methods on a set of 89 analogous protein pairs. The percentages denote the fraction of pairs for which a particular method yields a score. These cases are quantified in terms of the returned score (x-axis) and the coverage of the analogous region shared by the pairs (y-axis). The red vertical dashed lines correspond to the average score for homologous pairs.

A similar procedure was used to construct the “easy” benchmark set. As with the “hard” set, the ECOD was first clustered using MMSeqs2 with a 70% sequence identity cutoff to remove redundancy. The resulting sequences were then clustered again using a 50% sequence identity cutoff, and those from clusters containing less than seven members or containing members from more than one ECOD H-group were removed. Finally, for each H-group, only the largest MMSeqs2 cluster was retained. As a result, we obtained 1032 ECOD domains grouped into 103 clusters. In the “easy” benchmark, we used a stricter definition of positives and negatives, defining sequence pairs belonging to the same cluster as positives and the rest as negatives.

### Analogy benchmark

To assess the ability of the methods to distinguish homologous from analogous sequences, we used the MALISAM database (Cheng *et al*., 2008), which collects examples of protein pairs that share similar structural motifs that have evolved convergently. Eighty-nine such cases were obtained from the database website (http://prodata.swmed.edu/malisam/) and processed to define structurally alignable pairs of residues belonging to the analogous motifs. Each pair of sequences was aligned using pLM-BLAST in local and global modes, HHsearch, and EBA, and the resulting aligned pairs were compared with those from the MALISAM database. Based on these comparisons, the total coverage of the analogous motif and the score of the respective alignment were defined for each case (Figure 2B).

### Alignment quality benchmark

To evaluate the quality of the alignments generated by HHsearch, pLM-BLAST, EBA, and BLAST, we followed a procedure previously described in (Remmert *et al*., 2011). First, we extracted all pairs from the “hard” benchmark set (see above) for which all methods produced an alignment. Then, for each pair, we computed the structural alignment using the USalign tool (Zhang *et al*., 2022) and discarded those for which the TM score was below 0.5, resulting in 444 pairs. The structure-based sequence alignments provided by USalign defined the ground truth against which the alignments of each method were compared. Specifically, we considered the precision, i.e., the fraction of correctly aligned pairs out of all aligned pairs, and the sensitivity, i.e., the fraction of correctly aligned pairs out of all structurally alignable pairs. The precision and sensitivity values were averaged for each method and plotted (Figure 3). We also attempted to evaluate TM-Vec alignments, but for technical reasons were unable to run the DeepBlast model (Morton *et al*., 2020) required for alignment generation, but we expect it to perform with an accuracy comparable to other global alignment methods.

**Figure 3.**
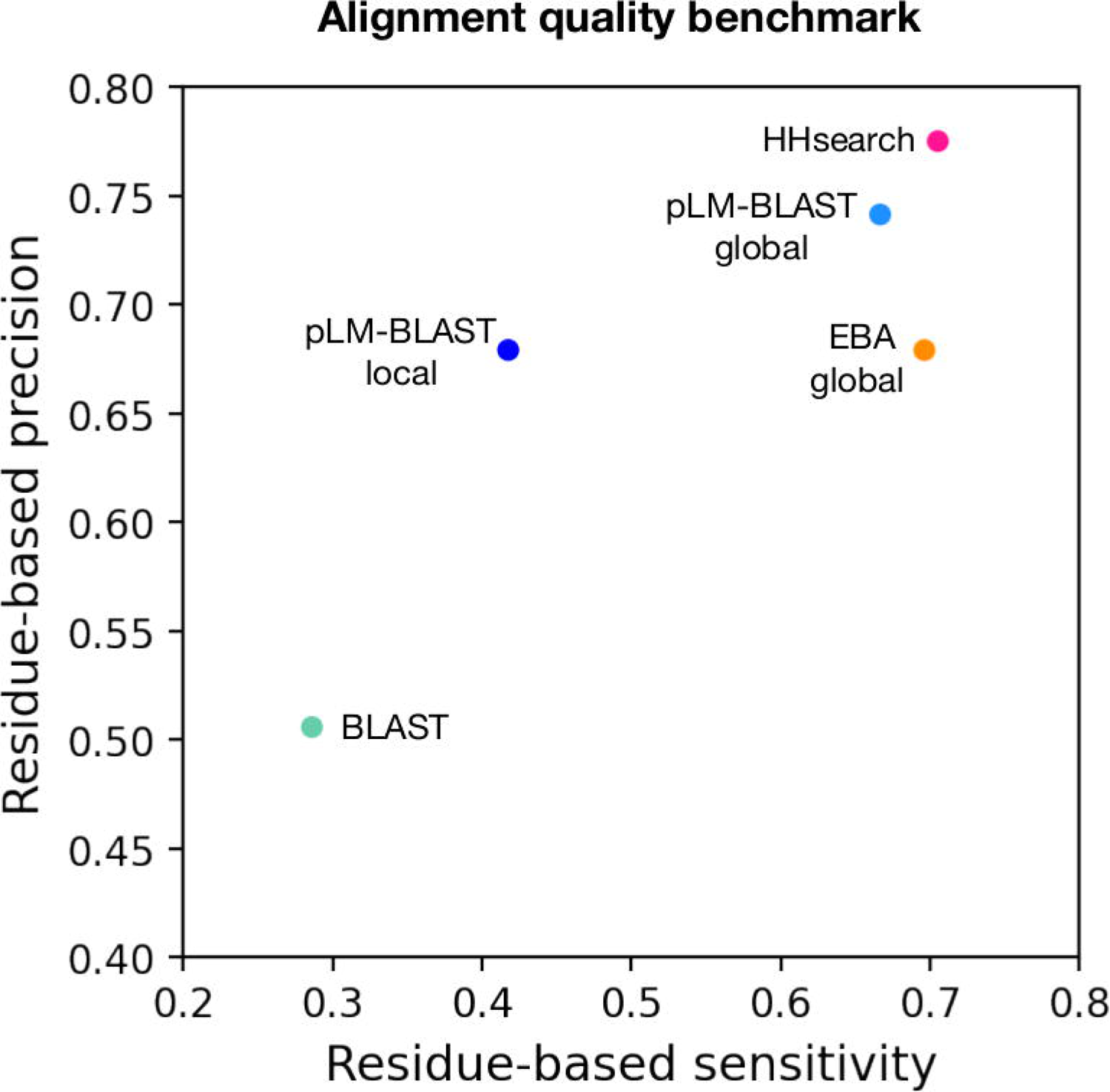
Alignment quality benchmark. Each point represents the average per-residue sensitivity (y-axis) and per-residue precision (x-axis) achieved in the reconstruction of 444 structure-based alignments by a specific method.

#### Speed Benchmark

To evaluate the speed of pLM-BLAST and other methods, we performed a benchmark using five query sequences (Supplementary File 3) of different lengths and folds to search the ECOD70 database, i.e., the ECOD database filtered to maximum 70% identity with MMseqs2. To generate the HHsearch database, we used HHblits run with 3 iterations on the UniRef30 database (UniRef database filtered to maximum 30% sequence identity), version 2022_02. The benchmark covered the time needed to prepare a given query (embedding calculation or HHblits search) and to search the database. pLM-BLAST was used with cosine similarity pre-filtering (see the next subsection) with the cut-off set at the 90th percentile. All calculations were performed for a fixed computer specification of a 6-core CPU and a single GPU. The benchmark was run 4 times for each method and the run times were averaged.

#### Cosine similarity pre-filtering

Typically, when dealing with embeddings, protein similarity is inferred as the cosine similarity between per-protein embeddings (global representations). This approach is very fast, but suffers from the loss of information stored in the per-residue embeddings (local representations) due to the averaging that occurs when the representation is converted to a fixed-size vector. This loss of information makes it impossible to capture local similarities and significantly reduces the ability to detect distant homology. With this in mind, we have developed a procedure that provides a trade-off between the use of global and local representations.

For a given pair of sequences (e.g., the query and one of the database sequences), per-residue embeddings are taken and treated as 2D dimensional images (Figure 1A), where one dimension is the sequence length and the other is the embedding size (1024 for ProtT5). We then convolve one embedding with another, where the first dimension of each convolution window is user-defined (30 by default) and the second dimension is fixed and equal to the size of the embedding dimension (1024 for ProtT5). This approach produces a 2D matrix in which each element represents the cosine similarity between each embedding slice of the two sequences. Finally, only if the resulting matrix has a value above the user-specified threshold, the corresponding sequences are passed to the actual pLM-BLAST method.

## Results and Discussion

### Homology detection

To assess the performance of pLM-BLAST in homology detection, we conducted a comparison with state-of-the-art methods such as BLAST and HHsearch, as well as other embedding-based methods (TM-Vec and EBA), using the Evolutionary Classification of Protein Domains (ECOD) database as a reference (Cheng *et al*., 2014). The ECOD database categorizes protein domains based on their evolutionary relationships and structural similarities and organizes them into families, topologies, homologous groups (H-groups), possibly homologous groups (X-groups), and architectures. To evaluate each method’s ability to reconstruct 300 selected ECOD H-groups, two metrics were used: the precision-recall curve and the ratio of true positives to false positives (as shown in Figure 2). For this study, we specifically curated the benchmark set to ensure that no pair of sequences exceeded 30% similarity, which led to a particularly challenging dataset that we refer to as the “hard” set.

In the “hard” benchmark, EBA performed best, followed by HHsearch and pLM-BLAST, both of which offer similar levels of accuracy; meanwhile, TM-Vec and BLAST underperformed (Figure 2). The unexpected discrepancy in performance between EBA, TM-Vec, and pLM-BLAST is noteworthy, given they all use the same representations derived from the ProtT5 language model. Unlike the better-performing methods, pLM-BLAST and EBA, which rely solely on unsupervised comparisons of raw representations, TM-Vec, the worst-performing method, uses ProtT5 representations as input for a specially trained end-to-end deep learning model to generate structural alignments. This suggests that the raw embeddings carry sufficient information for homology detection, whereas the structural similarity predicted by TM-Vec serves as a much less informative marker for homology.

The superior performance of EBA among ProtT5-based methods can be attributed to its utilization of an efficient signal enhancement procedure, which facilitates a more effective comparison of representations (Pantolini *et al*., 2022). It should also be noted that EBA provides only global alignments, which are expected to perform better than local alignments on a benchmark set consisting of domains, rather than full-length sequences. This relationship is evident in the performance comparison between pLM-BLAST running in local and global modes, with the latter consistently outperforming the former.

#### Analogy detection

Given the strong performance of pLM-BLAST and EBA in the distant homology detection benchmark, where they achieved accuracies comparable to HHsearch (Figure 2), we further tested them for their ability to distinguish homology from analogy (i.e., sequence similarity resulting from convergent evolution rather than common ancestry). To do this, we used 89 cases from the MALISAM database (Cheng *et al*., 2008), which collects examples of protein motifs that share similar structures due to analogous origins. HHsearch and pLM-BLAST’s local alignments covered the analogous regions in only 2% and 3% of the test cases, respectively, all with scores below those expected for clearly homologous pairs (Figure 2B). In contrast, pLM-BLAST and EBA global alignments covered analogous regions in 31% and 33% of cases, respectively, but also with scores below those expected for homologous sequences. These results suggest that embedding-based methods, much like HHsearch, are not misled by analogous signals, especially when local alignments are used.

#### Alignment quality

The benchmarks described above focused on the detection of homologous domain pairs based on the provided scores. However, an equally important aspect is the extent to which the generated alignments are correct at the per-residue level. To evaluate this, we used structure-based sequence alignments as a reference point against which the individual sequence-based alignments were compared. For each method, we calculated per-residue precision and sensitivity coefficients (see Methods for details). The results obtained indicate that pLM-BLAST global, EBA global, and HHsearch all provided very good alignments, with EBA global slightly less effective in terms of precision (Figure 3). The pLM-BLAST in local mode also provided good precision but lower sensitivity, indicating that its alignments are as precise as those of HHsearch and other embedding-based methods, which run in global mode but shorter, thus not covering all structurally alignable pairs. As we will illustrate later in the text, such short alignments may correspond to ancestral fragments (Alva *et al*., 2015; Kolodny, 2021) that are shared between different protein folds.

### Comparison of similar sequences

Above, we discussed pLM-BLAST and related methods in the context of detecting homology between highly divergent sequences that share less than 30% pairwise sequence identity. However, sequences sharing 50% or more identity are also routinely compared in bioinformatics tasks. Bearing this in mind, we considered an additional “easy” benchmark set, where homologous sequences share between 50% and 70% identity (see Methods for details). This benchmark was also based on domain sequences from the ECOD database, but it aimed to reconstruct clusters defined by MMseqs2 instead of ECOD H-groups.

The results obtained indicate that pLM-BLAST and all other benchmarked methods perform almost perfectly in this task (Figure 4A). Only at precision and recall rates greater than 95% can minor differences be observed, with BLAST outperforming the others and TM-Vec demonstrating the lowest accuracy. pLM-BLAST and EBA in global alignment mode perform equally well, closely followed by HHsearch and pLM-BLAST in local mode. The strong performance of pLM-BLAST, EBA, and TM-Vec in this test suggests that embedding-based methods are a viable option for comparing similar sequences and should be considered in the development of new tools for sequence searching and clustering, akin to DIAMOND (Buchfink et al., 2021) or MMseqs2 (Steinegger and Söding, 2017).

**Figure 4.**
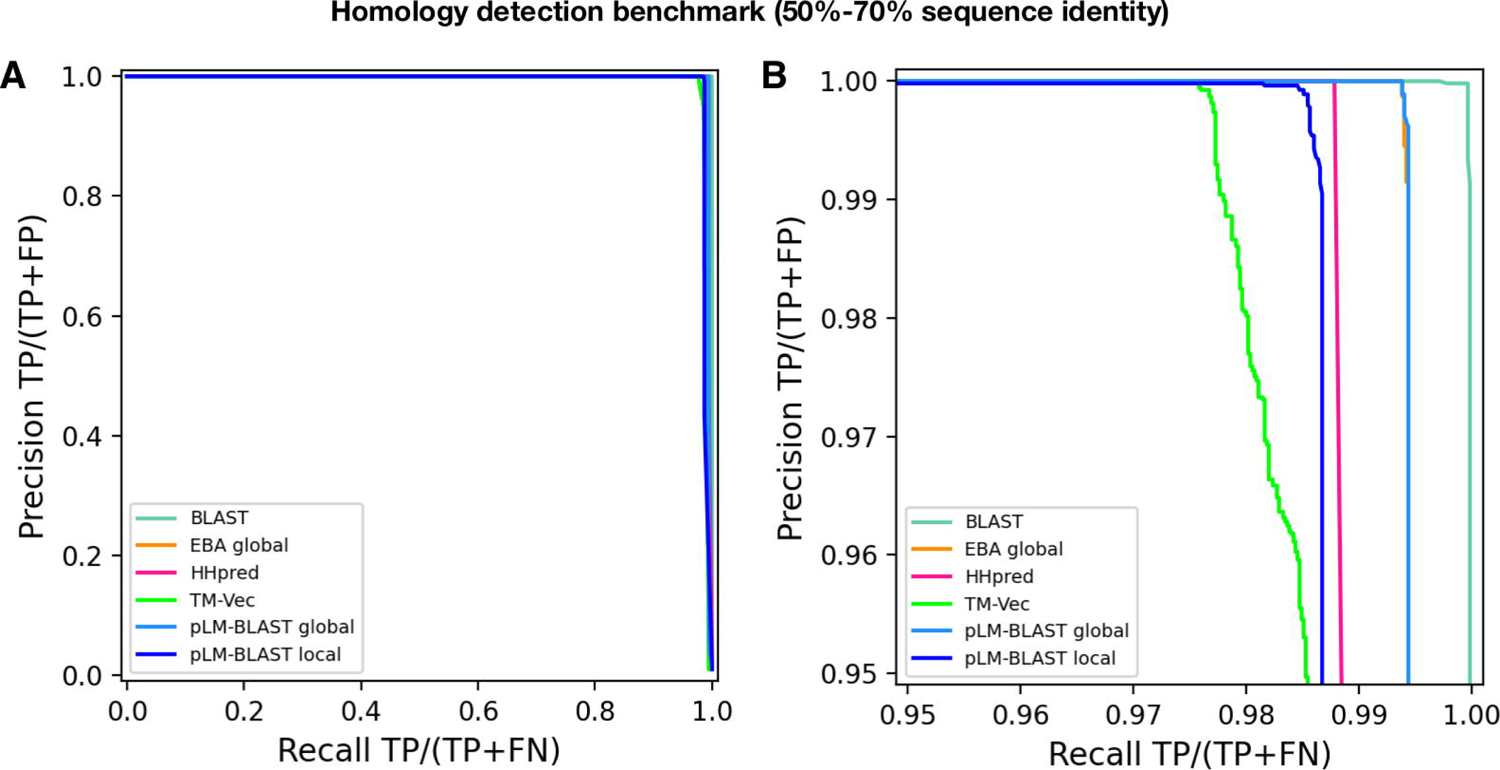
Performance of pLM-BLAST and other methods in classifying sequences with 50-70% identity. **(A)** Precision-recall curve. **(B)** The same precision-recall curve as in (A) is shown, but it is limited to the precision and recall ranges that are greater than 95%.

#### Speed benchmark

A critical aspect of large-scale analyses is computational speed. Bearing this in mind, we evaluated the speed of pLM-BLAST, EBA, HHsearch, and TM-Vec while searching the ECOD70 database (over 62, 000 records) using five example queries of varying lengths (Supplementary file 1).

The results reveal (Table 1) that TM-Vec is the fastest method, despite being the worst performer in the homology detection benchmark (Figures 2). pLM-BLAST, the second fastest method, is on average twice as fast as HHsearch and three times faster than EBA. The only case where HHsearch outperformed pLM-BLAST was a coiled-coil domain (query 2 in Table 1), where only a few homologous sequences could be detected. These results suggest that pLM-BLAST offers an optimal balance between accuracy and speed, with its performance being comparable to HHsearch (see Figures 2 and 3), yet faster, and offering new insights into protein evolution, as discussed later. EBA, which delivered the best performance in the homology detection benchmark, was also the slowest method. This is probably because it lacks the pre-filtering step employed by pLM-BLAST (see Methods).

**Table 1.** Speed comparison between pLM-BLAST and other methods. The runtimes are presented in seconds, and the length of each query sequence is provided in brackets.

| Method | Query 1<br>(269) | Query 2<br>(55) | Query 3<br>(379) | Query 4<br>(88) | Query 5<br>(132) |
| --- | --- | --- | --- | --- | --- |
| pLM-BLAST local | 2m 5s | 54s | 2m 30s | 1m 3s | 1m 27s |
| HHsearch | 5m 9s | 38s | 3m 4s | 4m 2s | 2m 57s |
| EBA | 7m 10s | 3m 15s | 7m 23s | 3m 46s | 5m 0s |
| TM-Vec | 15s | 15s | 15s | 15s | 15s |

### Potential to discover new homology relationships

Given the ability of pLM-BLAST to detect homology (Figure 2A), distinguish between homology and analogy (Figure 2B), and provide high-precision alignments (Figure 3), we speculated that some of the high-scoring connections between different H-groups of the same X-group, or even different X-groups, could reflect true homology.

In the benchmark of pLM-BLAST run in the local mode, the most number of connections between H-groups were found in the Alpha-beta plaits and Flavodoxin-like X-groups (see Supplementary File 2 for the full list of connected H-groups). Most of these intra-X-group matches were not detected by either HHsearch, which is expected since it was used to define homologous relationships in the ECOD database (Cheng *et al*., 2014), or EBA, which typically yielded low scores. We also detected connections between domains of different X-groups, often hinged on the presence of a subdomain-sized fragment (Alva *et al*., 2015; Kolodny, 2021) (Supplementary Table 2); for example, between the P-loop, Rossmann, and Flavodoxin folds (Longo *et al*., 2020; Kolodny *et al*., 2021). Among these cases, we found a fragment conserved between domains of two different sandwich folds, namely cupredoxin (ECOD H-group 3156.1) and immunoglobulin (11.1, Figure 5A). Despite a global structural similarity (DALI Z-score=2.7), the sequences of these two proteins could be aligned only over a short region corresponding to a beta-hairpin motif. This motif may represent an ancestral fragment that has been independently “decorated” with secondary structural elements, resulting in the two observed structures. In such a scenario, only the two beta-hairpin motifs would be truly homologous, while the overall structural similarity could be attributed to convergent evolution. Moreover, a homologous relationship between these two domains has been discussed previously (Gough and Chothia, 2004; Stevens, 2008), and a pLM-BLAST scan of the entire ECOD database using a cupredoxin domain as a query revealed additional connections to other Ig-like domains (Figure 5C). Among them, we identified structures from different ECOD H-groups of the immunoglobulin-like beta-sandwich X-group (also see Supplementary Table 2), suggesting homology in the region of the beta-hairpin motif. Interestingly, the matches also included a significantly different jelly-roll fold. Although the structural similarity between cupredoxin and jelly-roll folds is not as apparent as that of the of the cupredoxin and Ig-like folds, all share a Greek key motif in their core, lending credence to a hypothesis of a common origin.

**Figure 5.**
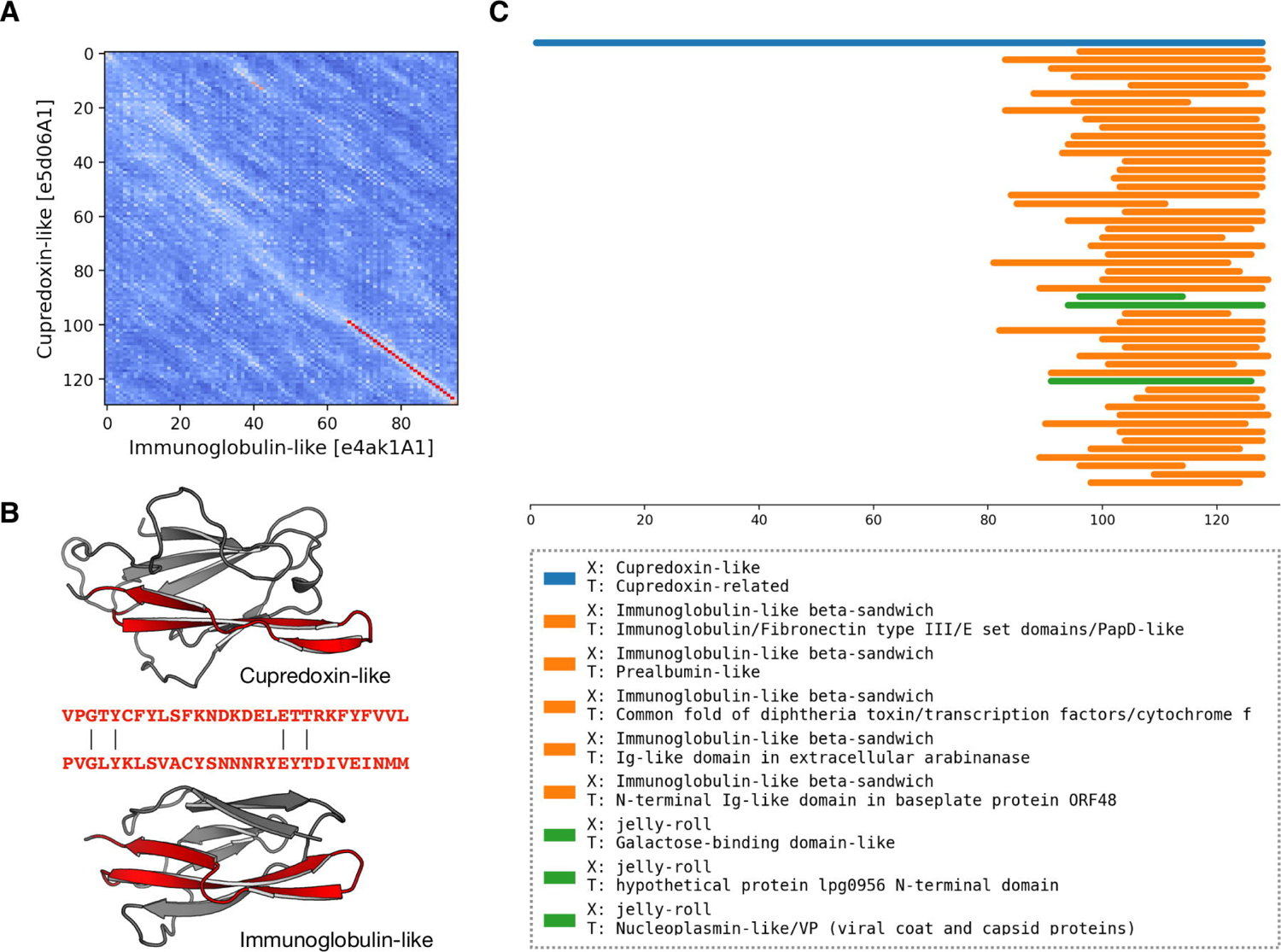
pLM-BLAST scan of the ECOD database using the sequence of the cupredoxin domain. **(A)** A substitution matrix illustrating the similarity between cupredoxin and immunoglobulin folds. The numbered axes indicate consecutive residues. The lighter the color, the higher the similarity between the sequences in a given region. The best local alignment is indicated by a red line. **(B)** Structures of cupredoxin and immunoglobulin folds, with the homologous region detected by pLM-BLAST highlighted in red. **(C)** Graphical representation of the scan results. Bars correspond to hits, color-coded according to their ECOD X-group membership.

In our benchmark, we used HHsearch’s homology predictions as a ground truth and reference point for pLM-BLAST and other methods. However, it is important to note that homology cannot be definitively proven, and the benchmark is essentially a comparison of one hypothesis against another. While HHsearch is a well-established method, and its predictions have provided crucial insights in numerous studies, our results suggest that embedding-based methods such as pLM-BLAST could potentially detect homology beyond the boundaries defined by HHsearch.

## Conclusions

In summary, pLM-BLAST is a sensitive tool for remote homology detection that is based on comparison of sequence representations obtained from the pLM ProtT5. It provides both global and local alignments, with the latter being crucial for detecting distant evolutionary relationships (Figure 5). Additionally, it is efficient in database searching (Table 1) due to the use of a pre-filtering step. pLM-BLAST is available both as a stand-alone package and as an easy-to-use web server within the MPI Bioinformatics Toolkit (Zimmermann *et al*., 2018), where it can be used to search precomputed databases. Currently, the primary limitations of pLM-BLAST are the maximum size of the target database ( as searching enormous, redundant databases containing millions of sequences would be computationally costly), and the fact that in local mode, it typically produces highly precise but shorter alignments than HHsearch. However, as demonstrated in Figure 5, these shorter alignments can indeed be meaningful. The significance of these alignments, often being sub-alignments of those provided by HHsearch (if any are provided), from an evolutionary perspective remains to be explored. We anticipate that pLM-BLAST will prove beneficial in studying protein function and evolution, including in the cases of singletons/orphan sequences and taxonomically-restricted genes (Barrera-Redondo *et al*., 2022), for which deep HMM profiles cannot be calculated.

## Acknowledgments

This work was supported by the First TEAM program of the Foundation for Polish Science, co-financed by the European Union under the European Regional Development Fund (grant POIR.04.04.00-00-5CF1/18-00). V.A. and S.D-H. were supported by institutional funds from the Max Planck Society.

