## Supplementary figures and images for "pLM-BLAST – distant homology detection based on direct comparison of sequence representations from protein language models"

### Supplementary Figure 1

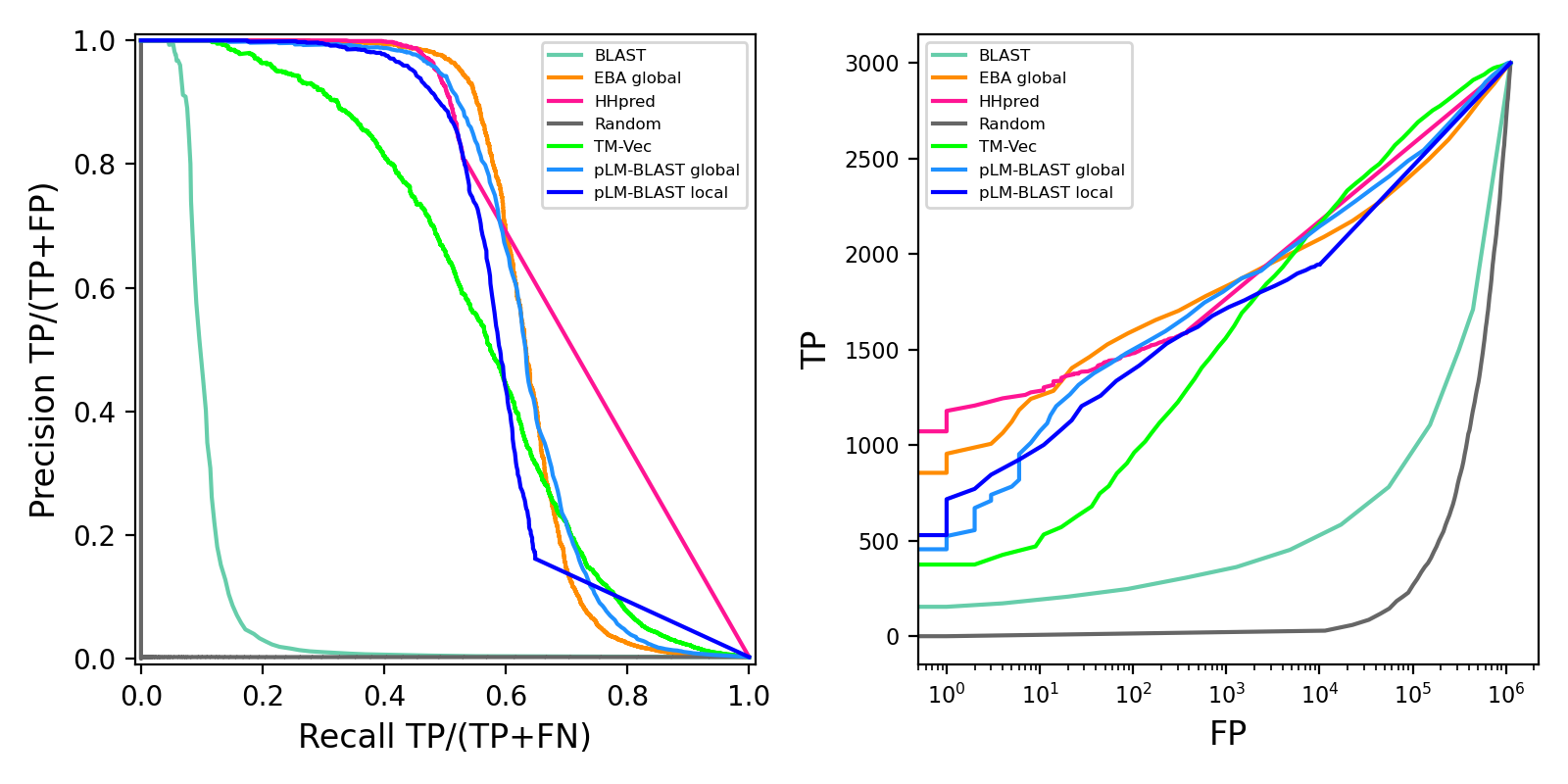
